# Specific domains of early parenting, their heritability and differential association with adolescent behavioural and emotional disorders and academic achievement

**DOI:** 10.1101/509513

**Authors:** Iryna Culpin, Marc H. Bornstein, Diane L. Putnick, Hannah Sallis, Ruby Lee, Miguel Cordero, Priya Rajyaguru, Katarzyna Kordas, Tim Cadman, Rebecca M. Pearson

## Abstract

**Background:** Variations in parenting across large populations have rarely been described. It also remains unclear which specific domains of parenting are important for which specific offspring developmental outcomes. The study describes different domains of early parenting behaviours, their genetic heritability, and then determines the extent to which specific domains of parenting are associated with later offspring outcomes.

**Methods:** Parenting behaviours (birth to 3 years) were extracted from self-reported questionnaires administered to 12,358 mothers from the UK-based birth cohort study, the Avon Longitudinal Study of Parents and Children, and modelled as a latent factor using Confirmatory Factor Analysis. Genetic heritability and correlations between parenting factors were estimated using wide complex trait analysis. Offspring emotional, behavioural and academic outcomes were assessed at age 16 years using the self-reported Short Mood and Feelings Questionnaire, the Development and Well-Being Assessment, and grades achieved in national English Language examination.

**Results:** Three parenting factors were derived: parental enjoyment, conflictual relationships and stimulation; all showed low genetic heritability. There was no evidence of associations between parenting factors and offspring depressed mood. Stimulation was associated with better English grades after controlling for maternal education (standardised β=0.058, p=0.007), and enjoyment was negatively associated with English grades (β=-0.082, p=0.002). Conflictual relationships were associated with higher risk of offspring behavioural disorders after controlling for behavioural problems at age 3 years (β=0.227, p=0.007). Higher enjoyment reduced the association between conflict and behavioural problems (interaction term β=0.113, p<0.001).

**Conclusions:** We found evidence for predictive specificity of early parenting domains for offspring outcomes in adolescence. Early stimulation, unlike enjoyment, promoted later educational achievement. Conflictual relationships were associated with greater risk of behavioural problems, buffered by increased enjoyment. These findings hold implications for parenting interventions, guiding their focus according to specificity of parenting domains and their long-term outcomes in children.

## Introduction

Variations in mother-child interactions and the quality of early parenting are associated with long-term child outcomes, including mental and physical health, socioemotional and cognitive development (Bornstein, 2015; Collins, Maccoby, Steinberg, Hetherington, & Bornstein, 2000). However, parenting is a complex construct. Parents not only nurture and protect children, they also guide them in understanding and expressing appropriate feelings and emotions, as well as educate and prepare them for adaptation to a wider range of life roles and contexts (Bornstein, 2015). They also deal with disciplining and conflicts as their children grow and take risks. Thus, parenting practices constitute a varied and demanding set of skills, and there is considerable variation in how adults esteem and execute different components of the caregiving repertoire. However, different aspects of the parenting experience across large populations of parents have rarely been described (Bornstein, 2016).

### Parenting and child outcomes

Apart from describing variations in parenting behaviours, it is important to understand the impact different parenting domains have on long-term child outcomes. Parenting practices are often cast along three main domains, warmth/love or enjoyment, control, discipline and conflict and stimulation (Bornstein, 2015). Warmth encompasses enjoyment, sensitivity and involvement (Pomerantz & Thompson, 2008), with higher levels of warm, sensitive and developmentally stimulating parenting being associated with decreased child negative reactivity (Bates et al., 2012) and greater academic achievement (Boivin & Bierman, 2013). On the contrary, parent-child relationships characterized by relatively low levels of warmth and affective enjoyment are associated with offspring emotional (Rudolph et al., 2000) and behavioural (Olson, Bates, Sandy, & Lanthier, 2000) problems in adolescence. Parental control ranges from monitoring to harsh discipline (Berger & Langton, 2011), with higher levels of conflicts within the parent-child relationship and harsh discipline being associated with offspring behavioural problems in adolescence (Rajyaguru, Moran, Cordero, & Pearson, 2018). Stimulation, defined as parental activities to promote learning (Lugo‐Gil & Tamis‐LeMonda), has been found to predict offspring cognitive abilities (Landry, Smith, & Swank, 2006) and academic achievement (Gottfried, Marcoulides, Gottfried, & Oliver, 2009).

### Importance of specificity

Although links between parenting and child outcomes are well-documented (Bornstein, 2015; Collins et al., 2000), the importance of specific aspects of parenting for particular child outcomes has rarely been studied. Parenting interventions can be complex and taking a ‘one size fits all’ approach is often ineffective, with many universal efforts failing to show evidence of positive effects across all outcomes (Triple P-Positive Parenting Program; Marryat, Thompson, & Wilson, 2017; The Nurse-Family Partnership; Olds, Hill, O’Brien, Racine, & Moritz, 2006). Thus, it is essential to establish links between specific parenting domains and specific child outcomes to design targeted parenting interventions. Here we examine the extent to which specific parenting domains are associated with offspring outcomes. We hypothesise that enjoyment/warmth will be associated with emotional outcomes, whilst conflict with behvaioural, and stimulation with academic offpring outcomes.

### Genetic basis of different domains of parenting

Variations in different components of parenting will be driven by both genetic and environmental factors (Bornstein, 2016). This information is important to understand the extent to which parenting can be modified. Meta-analysis of previous research based on twin and adoptive studies indicates a moderate genetic basis (23%-40%) across most parenting phenotypes (Khlar & Burt, 2013), with some evidence for variation in genetic influence depending on the parenting components measured. For instance, parental genetic factors explained less of the ‘negative’ aspects of parenting such as conflict with children than the ‘positive’ aspects such as warmth and enjoyment (Klahr & Burt, 2013). However, these studies have used twin studies to estimate Heritability (h^2^): the proportion of variation in a phenotype that can be attributed to genetic differences for the particular context and timepoint). Twin models, however often over estimate heritability (Sallis, Davey Smith, & Munafo, 2018). An alternative approach is to use molecular genetic data and estimate the heritability captured by single nucleotide polymorphisms (SNPs) included on genotyping platforms (Yang, Lee, Goddard, & Visscher, 2011). This has not been applied to the heritability of parenting before. Here we describe different components of parenting experiences and estimate SNP based h^2^ from maternal molecular genetic data.

### Current study

In the current study, we address the limitations of previous research by describing the different domains of self-reported parenting in the first 3 years of life and estimating the extent to which these domains are associated with emotional and behavioural disorders and academic achievement in offspring at age 16 years using data from a large UK-based birth cohort study, the Avon Longitudinal Study of Parents and Children (ALSPAC). The unique richness of the ALSPAC data provides a rare opportunity to control for early measures of child behavioural problems that may affect parenting, thus, controlling for reverse causality. We also utilise molecular genetic data to estimate the variance explained by genetic factors and to examine shared genetic architecture across factors.

## Methods

### Sample

The sample comprised participants from the Avon Longitudinal Study of Parents and Children (ALSPAC). During Phase I enrolment, 14,541 pregnant mothers residing in the former Avon Health Authority in the south-west of England with expected dates of delivery between 1 April 1991 and 31 December 1992 were recruited. When the oldest children were approximately 7 years of age, an attempt was made to bolster the initial sample with eligible cases who had failed to join the study originally. The total sample size for analyses using data after the age of 7 years is 15,247 pregnancies, of which 14,899 were alive at 1 year of age. Our sample comprised 12,358 mothers with at least one parenting item. Ethical approval and informed consent for the study was obtained from the ALSPAC Ethics and Law Committee and the Local Research Ethics Committees. Informed consent for the use of data collected via questionnaires was obtained from participants following the recommendations of the ALSPAC Ethics and Law Committee at the time. Detailed information about the cohort has been collected since early pregnancy, including regular self-completion questionnaires from mothers and children. Information about ALSPAC is available at www.bristol.ac.uk/alspac/, including a searchable data dictionary (http://www.bris.ac.uk/alspac/researchers/data-access/data-dictionary/). Further details on the cohort profile, representativeness and phases of recruitment are described in two cohort-profile papers (Boyd et al., 2013; Fraser et al., 2012).

## Measures

### Development of Parenting Factors

#### Process of item selection

Full details of item section and development of parenting factors are provided in Supplementary Material. In summary, potential items were extracted from self-reported questionnaires administered from pregnancy to age 3 years capturing parenting behaviour, attitudes and knowledge. Items categorised as parental enjoyment, conflictual relationships, and stimulation and teaching (based on parenting taxonomies; Maccoby & Martin, 1983) were extracted and entered into separate single-factor Confirmatory Factor Analysis (CFA) models. We focused on ages 0-3 years to capture a period of time most mothers spend with their children prior to the commencement of nursery school.

### Adolescent outcomes

#### Depressed mood

The Short Mood and Feelings Questionnaire (SMFQ; Angold, Erkanli, Silberg, Eaves, & Costello, 2002) was administered at age 16 years via postal questionnaires. It consists of 13 items relating to low mood during the past two weeks, each with scores of 0 to 2. Individual items are summed, producing a total score ranging from 0 to 26, which was dichotomised to classify individuals as depressed *versus* not-depressed using a cut-off point of ≥ 11. This cut-off point has been shown to have high sensitivity and specificity (Thapar & McGuffin, 1998) and has been applied in previous studies based on community samples (Patton et al., 2008).

#### Behavioural disorders

Behavioural disorders were assessed using parent and child versions of the Development and Well-Being Assessment (DAWBA; Goodman et al., 2000). The semi-structured interview comprises questions about a range of symptoms relevant to childhood psychiatric disorders. At age 15 years, the parent-completed DAWBA was used to assess symptoms of Disruptive Behaviour Disorder (DBD), Oppositional Defiant Disorder (ODD), and Attention Deficit Hyperactivity Disorder (ADHD) over the past 6 months, or conduct disorder over the past year. Children are not asked in detail about behavioural disorders due to possible bias in reporting these conditions (Loeber, Green, Lahey, & Stouthamer-Loeber, 1991). Child-reported versions of the DAWBA were used to assess symptoms of Major Depressive Disorder (MDD) and Generalised Anxiety Disorder (GAD) over the past 6 months. Binary variables were derived to characterise diagnosis of emotional and behavioural disorders (*versus* no diagnosis).

#### Educational achievement

General Certificate of Secondary Education (GCSE) grades achieved in English Language at age 16 years were extracted from external educational records and with consent linked to ALSPAC identification numbers. A binary variable was created to represent either failing to achieve an A*-C in English (coded as 1) or achieving A*-C (coded as 0), which is an essential qualification in the UK, as all other outcomes were negative we coded in this direction. We focused on the GCSE grades in English only to avoid multiple comparisons with a number of GCSE grades. In addition, grades in English are the most relevant outcome to the parenting domain of stimulation and teaching (mostly composed of language-related items: e.g., story telling, song singing).

#### Confounding variables

Parental and child characteristics identified in previous studies as being associated with parenting and child outcomes were accounted for in the model. These included highest maternal educational attainment (minimal education or none, compulsory secondary level (up to age 16 years), non-compulsory secondary level (up to age 18 years) *versus* university level education), maternal antenatal depression (18 and 32 weeks’ gestation) assessed using the Edinburgh Postnatal Depression Scale (EPDS; Cox, Holden, & Sagovsky, 1987), maternal age (in years), child gender (male *versus* female) and early behavioural problems assessed at age 3 years through maternal reports using the total problems scale of the Rutter revised preschool questionnaire (Elander & Rutter, 1996).

### Analysis

Models were estimated using Structural Equation Modelling (SEM) in M*plus* v.7 (Muthén & Muthén, 2015). Full Information Maximum Likelihood (FIML; Arbuckle, 1996) estimator was used to account for the missing data. FIML renders unbiased and more efficient estimates under Missing-At-Random (MAR) missing data conditions (Enders & Bandalos, 2001). A model was considered to have a good fit if the Root Mean Square Error of Approximation (RMSEA) was ≤ 0.06, and the Comparative Fit Index (CFI) and Tucker-Lewis Index (TLI) cut-off values was close to 0.95 (Hu & Bentler, 1998). The chi-square test of overall fit is sensitive to model misspecification in instances when sample size is large (Kline, 2005), thus, we gave greater weight to the incremental fit indices (Hu & Bentler, 1998).

#### Latent factor model

Full details of latent factor model derivation, including the flow chart of items included into the CFA (Figure 1S) and derived factors and factor loadings (Table S1), are presented in the Supplementary Material. In brief, items that were both theoretically assigned and showed standardised loadings >0.15 on the relevant dimension were entered into a combined model using Confirmatory Factor Analyses (CFA) with a robust weighted least square (WLSMV) estimator to model categorical data (Brown, 2006).

#### Estimating heritability of each of the 3 parenting factors and genetic correlation between parenting factors

Analyses to estimate heritability and genetic correlations are described in more detail in the Supplementart Material. In summary, we first calculated estimates of SNP-based heritability (h^2^ _SNP_) for each parenting factor using the restricted maximum likelihood (REML) method implemented within the genome-wide complex trait analysis (GCTA) software (Yang, Lee, Goddard, & Visscher, 2011). Second, we used a bivariate REML approach to estimate the genetic correlation between each of the parenting factors with each other to investigate shared genetic architecture across each of the parenting factors. Any overlap here could be due to pleiotropy (genetic effects on multiple traits), shared biological mechanisms between domains, or a causal relation from one domain to another.

## Results

Associations between parenting factors and child and parental confounders are presented in Table S2, Supplementary Material. Characteristics of the sample by the completeness of data are presented in Table S3, Supplementary Material.

### Final latent parenting factors

A model using CFA to fit the follwing 3 factors with the following items showed good model fit. The RMSEA (0.024, 95% CI 0.024 to 0.025) and the CFI (0.92) indicated that the measurement model fit the data well, supporting the adequacy of the model for tests of structural paths. There were relatively high correlations between parenting factors (Figure 1), however, a second order or bi-factor model did not converge or fit the data well.

**Figure 1.**
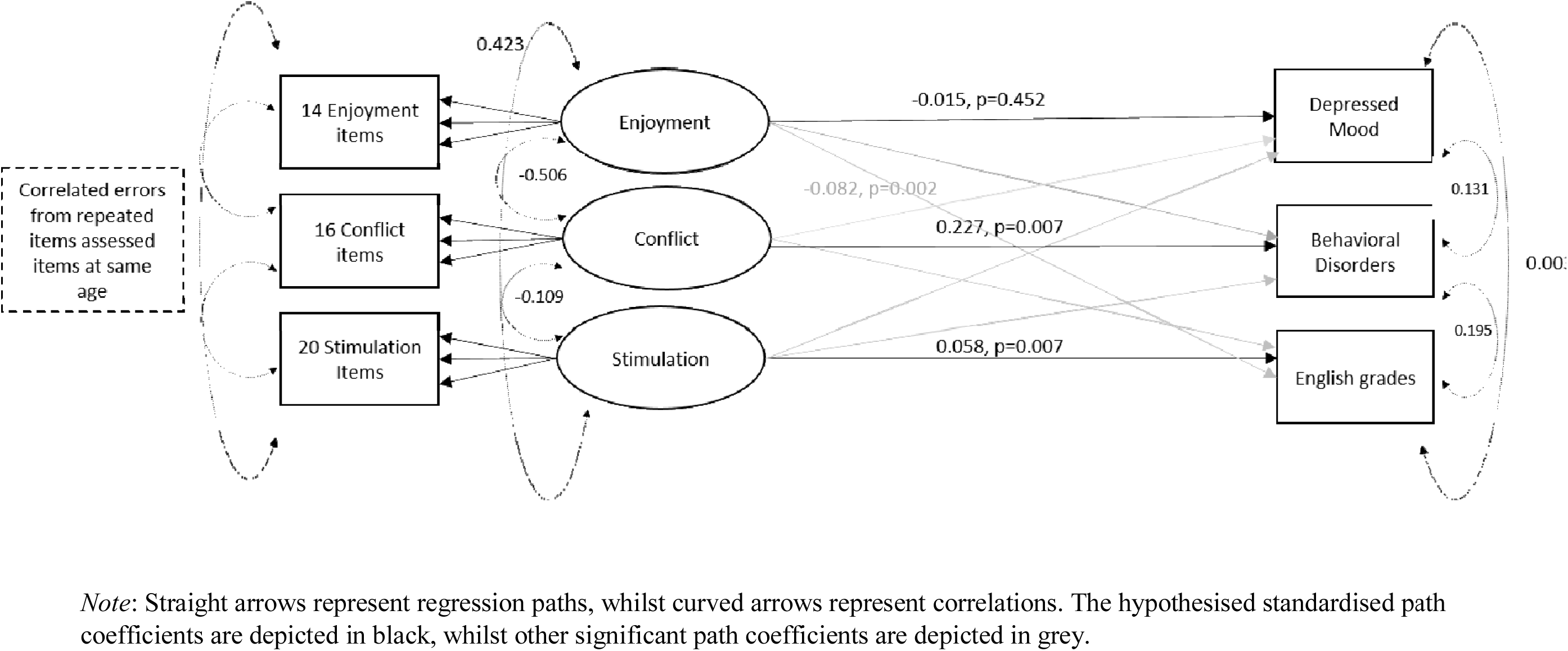
Latent factor model representing associations between parenting factors and adolescent outcomes following adjustments for parental (maternal age, educational attainment, prenatal depression) and child (gender and early child behavioural problems) confounders. Analyses conducted on all available data for each estimate using WLSMV defaults in Mplus (n=12,358).

#### Factor 1: Parental Enjoyment

Parental enjoyment contains 14 items relating to enjoyment of the child from ages 4 weeks to 3 years 11 months (e.g., ‘I really enjoy my baby’, ‘Having a baby has made me feel more fulfilled’) as well as items relating to frequency of cuddling and playing with the child. Initially, items relating to feelings of irritation with the child (e.g., ‘This child gets on my nerves’) were included, however, in the final model they loaded better on the factor encapsulating conflictual mother-child relationship. The internal consistency of parental enjoyment is α=0.82, with the summed items forming a normally distributed scale. Higher factor scores represent less parental enjoyement.

#### Factor 2: Conflictual Relationships

Conflictual relationships contains 16 items relating to conflict, harsh discipline and irritation with the child (e.g., frequency of arguments, ‘battle of wills’, smacking and shouting) from ages 1 year 6 months to 3 years 11 months. At age 1 year 6 months, a substantial proportion of mothers reported having battles of wills (37%) and frequent conflict (21%) with their children. In addition, 24% of mothers reported having smacked their children sometimes during tantrums, whilst 58% of mothers reported having shouted at their child. At age 3 years 11 months, 17% of mothers reported that the child gets on their nerves. The internal consistency of conflictual relationships is α=0.75. Higher factor scores signify more conflictual relationships.

#### Factor 3: Stimulation and Teaching

Stimulation and teaching contains 20 items relating to the frequency of engagement in teaching and stimulating activities from ages 6 months to 3 years 6 months (e.g., naming parts of the body, colours, numbers, singing to and talking with the child). At age 6 months, a substantial proportion of mothers reported often teaching (37%) and talking (30%) to their child, whilst 62% reported always talking to their child when doing household activities. At age 1 year 6 months, a majority of mothers reported that they teach their child the alphabet (70%), songs (7%), and politeness (4%). At age 2 years, 83% of mothers reported that they take their child to the park or playground at least once a week. The internal consistency for stimulation and teaching is α=0.75. Higher factor scores represent less stimulation and teaching.

### Associations between parenting factors and offspring behavioural disorders, depressive symptoms and academic achievement at 16 years

Latent parenting factors were regressed onto offspring depressive symptoms, behavioural disorders and academic achievement at age 16 years in the same model, leading to mutually adjusted associations between each parenting factor and each adolescent outcome. The model was adjusted for a number of possible parental (maternal age, educational attainment, depression) and child (gender and early behavioural problems) confounders. Given the complexity of the model, interaction terms were derived from saved factor scores for each latent factor to investigate interactions between parenting factors. The 3 interaction terms between continuous scores (stimulation*enjoyment; stimulation*conflict and conflict*enjoyment) were regressed onto each of the outcomes, with parenting factor scores also entered into the model.

Standardised path coefficients (β) of the associations between parenting factors and offspring emotional, behavioural and academic outcomes are presented in Table S4, Supplementary Material. There was evidence for a strong association between conflictual relationships and offspring behavioural disorders, but not depressive symptoms or educational achievement, at age 16 years. This longitudinal association remained independent following adjustment for parental reports of behavioural problems at age 3 years (β=0.227, p=0.007, 95% CI 0.062 to 0.391). There was no evidence for an independent association between early childhood behavioural problems and adolescent behavioural disorders in the mutually adjusted model. There was also evidence for an interaction between conflictual relationships and enjoyment with offspring behavioural disorders at age 16 years (interaction term β=0.113, p<0.001; Figure S2, Supplementary Material, represents percentage of offspring with CD or ODD diagnosis according to patterns of parental conflict and enjoyment). There was no evidence for an independent association between any of the parenting factors or their interaction and offspring depressive mood at age 16 years.

Early teaching and stimulation activities were associated with better GCSE grades in English Language at age 16 years after controlling for maternal education, although the effect sizes were relatively small (β=0.058, p=0.007, 95% CI 0.015 to 0.101). There was evidence for a negative association between parental enjoyment and English grades, independent of teaching and stimulation (β=-0.082, p=0.002, 95% CI −0.135 to −0.030). However, there was no evidence for an interaction between parental enjoyment and stimulation. Results were comparable when using complete case sample on all variables (n=2,694; Results S1, Supplementary Material).

### Heritability estimates of and genetic correlations between parenting factors

Estimates of h^2^_SNP_ were estimated for each of the parenting factors. Although effect sizes were small for each of the factors (suggesting that only a small proportion of the variation in each phenotype is due to genetic variation), larger estimates were observed for both conflictual relationships (h^2^_SNP_=0.055, se=0.05) and stimulation (h^2^_SNP_=0.036, se=0.05) than enjoyment (h^2^_SNP_ =0.002, se=0.05), for which h^2^_SNP_ was close to the null. However, these analyses were underpowered, and estimates should be interpreted with caution.

Estimates of genetic correlation suggested that the SNP effects for conflictual relationships and low enjoyment act in the same direction (r_G_=1.00, se=7.53), while SNP effects between conflictual relationships (r_G_=-0.646, se=1.12) and low enjoyment (r_G_=-1.00, se=9.09) with stimulation are negative.

## Discussion

The current study describes three different domains of self-reported early parenting behaviour, estimates the extent to which the parenting domains are heritable, and provides longitudinal evidence that links specific domains of parenting with specific offspring outcomes in adolescence, whilst estimating the proportion of variation in these domains that may be attributed to genetic factors. Estimates of heritability for each factor were small. However, given the small sample size, the confidence intervals do include larger estimates, and, therefore, we cannot rule out large estimates from this study alone.

Our findings indicate that conflictual parenting in the first 3 years of life is strongly associated with offspring behavioural disorders at age 16 years. This finding is consistent with previous research implicating harsh parental discipline in offspring behavioural problems (Brenner & Fox, 1998). This effect may be partly explained by reverse causality, whereby children who exhibit difficult behaviour contribute to a conflictual parent-child relationship. Indeed, a strong association emerged between parent-child conflict and child conduct problems at age 3 years. However, the longitudinal association between conflictual relationships in the parent-child dyad and adolescent behavioural disorders remained after adjustment for early childhood behavioural problems. In addition, there was no association between early childhood behavioural problems and adolescent behavioural disorders, suggesting that a conflictual parent-child relationship is a better predictor of conduct disorders in adolescence than early behavioural problems.

We found an interaction between conflictual parent-child relationship and enjoyment on offspring behavioural disorders at age 16 years, suggesting a possible ‘buffering’ effect of high parental enjoyment on the negative effect of conflictual and harsh parenting and associated behavioural disorders in adolescence. The mechanisms that underlie such ‘buffering’ by enjoyment remain unclear. It may be that the type of conflict encountered by parents and children reporting *both* high conflict and enjoyment is different from those who experience conflictual relationship without enjoymenet in other areas of the relationship. For instance, mothers who report high levels of conflict and enjoyment may be more emotionally expressive and have conflicts that, although frequent, are more quickly resolved. High levels of enjoyment may also facilitate a positive emotional environment where arguments and conflict are regularly resolved, and parents and children share positive feelings that further enhance positive parenting and optimal child development (Forgatch & DeGarmo, 1998).

We found no evidence for an independent association between any parenting factor or their interaction and offspring depressive mood at age 16 years. This is not to say that early parenting is inconsequential for adolescent depressive mood; rather, the 3 parenting factors we derived may not capture particular aspects of parenting related to offspring emotional development. For instance, it has been suggested that parental emotional scaffolding and regulation specifically in response to distress, as well as emotional availability, may be important for child emotional functioning (Katz, Maliken, & Stettler, 2012). This domain was not specifically captured here.

Unsurprisingly, early teaching and stimulation activities (e.g., reading, storytelling) were associated with better GCSE grades in English Language at age 16 years, even though the effect sizes were relatively small. Conversely, parental enjoyment was negatively associated with English grades at the end of school. The lack of interaction between parental enjoyment and stimulation suggests that, even in the context of high stimulation, enjoyment is still negatively associated with offspring academic achievement. A parent’s focus on low demandingness and letting children enjoy themselves, rather than enforcing learning (Baumrind, 1991), may eventuate in lower achievement. It should be noted, however, that our findings point to the importance of enjoyment for other offspring outcomes, such as its possible protective role in the association between conflictual parent-child relationship and adolescent behavioural disorders.

The strengths of this study include the large sample size, the long-term follow-up, the availability of repeated measures on parenting behaviour across early childhood as well as rich data on confounders and a longitudinal design that enabled examination of associations between early parenting and offspring emotional, behavioural, and academic adjustment in adolescence. Although it is likely that genetic analyses were underpowered, we were able to utilize molecular genetic data to estimate the proportion of variation in the parenting domains that could be attributed to genetic factors. We also accounted for reverse causality through adjustment for early child behavioural problems.

The findings need to be interpreted in light of several limitations. First, despite the population-based study design, it was impossible to rule out selection bias in relation to baseline recruitment or attrition in the sample over time. We attempted to address this concern by controlling for factors known to predict attrition in ALSPAC (e.g., parental education and psychopathology) and by using FIML estimator in M*plus* to account for missing data (Arbuckle, 1996). Second, we relied on parental reports of parenting behaviour, which may be subject to measurement error. However, measurement error is found in all measures of behaviour, including self-report and directly observed measures (Sessa, Avenevoli, & Steinberg, 2001). Arguably, for the dimensions of parenting under investigation, which involve rare events, such as harsh discipline and internal feelings of love or irritation, parental report is the most appropriate measure (Podsakoff, MacKenzie, Lee, & Podsakoff, 2003). Direct observations of parent-child interactions do not capture such events, whilst it is not possible to collect child-reported parenting between birth and age 3 years. In the present study, however, parenting factors were modelled using a latent variables approach, which explicitly accounts for measurement error by only modelling variance which is shared across items and separating this from specific variance likely reflecting error (Grewal, Cote, & Baumgartner, 2004).

## Conclusions

This study shows that different domains of parenting are important for different offspring outcomes. Early conflictual parent-child relationships in the context of low parental enjoyment are a strong predictor of offspring behavioural disorders, whilst early teaching and stimulation are important for subsequent academic achievement in English. This finding implies that strategies to reduce conflicts but also increase parental enjoyment of the child may be one avenue to reduce the risk of later offspring behavioural problems in families with conflictual parent-child relationships. Furthermore, encouraging early stimulation activities, such as more frequent reading and learning of simple concepts (e.g., alphabet), likely promotes subsequent offspring academic achievement.

## Supporting information

Supplemental Results and Tables

## Acknowledgements

We are extremely grateful to all the families who took part in this study, the midwives for their help in recruiting them, and the whole ALSPAC team, which includes interviewers, computer and laboratory technicians, clerical workers, research scientists, volunteers, managers, receptionists and nurses. The UK Medical Research Council and Wellcome (Grant ref: 102215/2/13/2) and the University of Bristol provide core support for ALSPAC. A comprehensive list of grants funding is available on the ALSPAC website (http://www.bristol.ac.uk/alspac/external/documents/grant-acknowledgements.pdf). This research was specifically funded by the European Research Commission awarded to Dr Pearson (Grant ref: 758813 MHINT). Professor Bornstein was funded by the Intramural Research Program of the NIH/NICHD, USA, and an International Research Fellowship in collaboration with the Centre for the Evaluation of Development Policies (EDePO) at the Institute for Fiscal Studies (IFS), London, UK, funded by the European Research Council (ERC) under the Horizon 2020 research and innovation programme (grant agreement No 695300-HKADeC-ERC-2015-AdG). Dr Cadman received funding from the Uropean Union’s Horizon 2020 research and innovation programme under grant agreement N: 733206, LIFE-CYCLE project. The UK Medical Research Council supports the MRC Integrative Epidemiology Unit (MC_UU_12013/4). This study was also supported by the NIHR Biomedical Research Centre at the University Hospitals Bristol NHS Foundation Trust and the University of Bristol. This publication is the work of the authors who will serve as guarantors for the contents of this paper. The views expressed in this publication are those of the author(s) and not necessarily those of the NHS, the National Institute for Health Research or the Department of Health.

## Key points

1. Parenting is a complex construct and variations across large populations have not been adequately described.
2. Specific aspects of early parenting are important for specific offspring outcomes in adolescence.
3. Three dimensions of parenting emerged in this large-population study: enjoyment, conflictual relationships and stimulation/teaching. All showed low genetic heritability and correlations between domains of parenting.
4. We found evidence for specificity of early parenting and differential effects on offspring emotional, behavioural, and educational outcomes in adolescence.
5. These findings hold important implications for the design of future parenting interventions guiding their focus according to parenting factors and offspring long-term outcomes.

## Abbreviations

ADHD: Attention Deficit Hyperactivity Disorder
DBD: Disruptive Behaviour Disorder
GAD: Generalised Anxiety Disorder
CD: conduct disorder
ODD: Oppositional Defiant Disorder
MDD: Major Depressive Disorder
GCSE: General Certificate of Secondary Education

## References

1. Angold, A., Erkanli, A., Silberg, J., Eaves, L., & Costello, E. J. (2002). Depression scale scores in 8–17-year-olds: effects of age and gender. Journal of Child Psychology and Psychiatry, 43, 1052–1063.

2. Arbuckle, J. L. (1996). Full information estimation in the presence of incomplete data. Advanced structural equation modelling: issues and techniques, 243, 277–365.

3. Bates, J. E., Schermerhorn, A. C., & Petersen, I. T. (2012). Temperament and parenting in developmental perspective. Handbook of temperament, 425–441.

4. Baumrind, D. (1991). The influence of parenting style on adolescent competence and substance use. The Journal of Early Adolescence, 11, 56–95.

5. Berger, L. M., & Langton, C. E. (2011). Young disadvantaged men as fathers. The Annals of the American Academy of Political and Social Science, 635, 56–75.

6. Boivin, M., & Bierman, K. L (Eds.). (2013). Promoting school readiness and early learning: implications of developmental research for practice. Guilford Publications: New York, NY.

7. Bornstein, M. H. Determinants of parenting. (2016). In D. Cicchetti (Ed.), Developmental Psychopathology: Risk, Resilience, and Intervention (3rd ed., Vol. 4, pp. 180–270). Hoboken, NJ: Wiley.

8. Bornstein, M. H. (2015). Children’s Parents. In M. H. Bornstein & T. Leventhal (Eds.), Ecological settings and processes in developmental systems. Handbook of child psychology and developmental science (Vol.4, 7^th^ ed.). Hobokeno, NJ: Wiley.

9. Boyd, A., Golding, J., Macleod, J., Lawlor, D. A., Fraser, A., Henderson, J., Molloy, L., Ness, A., Ring, S., & Davey Smith, G. (2013). Cohort profile: the ‘Children of the 90s’-the index offspring of the Avon Longitudinal Study of Parents and Children. International Journal of Epidemiology, 42, 111–127.

10. Brenner, V., & Fox, R. A. (1998). Parental discipline and behavior problems in young children. The Journal of Genetic Psychology, 159, 251–256.

11. Brown, T. A., & Moore, M. T. (2012). Confirmatory factor analysis. Handbook of structural equation modeling, 361–379.

12. Collins, W. A., Maccoby, E. E., Steinberg, L., Hetherington, E. M., & Bornstein, M. H. (2000). Contemporary research on parenting: The case for nature and nurture. American Psychologist, 55, 218–232.

13. Cox, J. L., Holden, J. M., & Sagovsky, R. (1987). Detection of postnatal depression: development of the 10-item Edinburgh Postnatal Depression Scale. The British Journal of Psychiatry, 150, 782–786.

14. Elander, J., & Rutter, M. (1996). Use and development of the Rutter parents’ and teachers’ scales. International Journal of Methods in Psychiatric Research, 6, 63–78.

15. Enders, C. K., & Bandalos, D. L. (2001). The relative performance of full information maximum likelihood estimation for missing data in structural equation models. Structural Equation Modelling, 8, 430–457.

16. Forgatch, M. S., & DeGarmo, D. S. (1998). Two faces of Janus: Cohesion and conflict. In Eds. Cox, M. J., Brooks-Gunn, J. Conflict and cohesion in families (pp. 181-198). Routledge.

17. Fraser, A., Macdonald-Wallis, C., Tilling, K., Boyd, A., Golding, J., Davey Smith, G., Henderson, J., Molloy, L., Ness, A., Ring, S., Nelson, S. M., & Lawlor, D. A. (2012). Cohort profile: the Avon Longitudinal Study of Parents and Children: ALSPAC mothers cohort. International Journal of Epidemiology, 42, 97–110.

18. Goodman, R., Ford, T., Richards, H., Gatward, R., & Meltzer, H. (2000). The Development and Well-Being Assessment: description and initial validation of an integrated assessment of child and adolescent psychopathology. The Journal of Child Psychology and Psychiatry and Allied Disciplines, 41, 645–655.

19. Gottfried, A. E., Marcoulides, G. A., Gottfried, A. W., & Oliver, P. H. (2009). A latent curve model of parental motivational practices and developmental decline in math and science academic intrinsic motivation. Journal of Educational Psychology, 101, 729–739.

20. Grewal, R., Cote, J. A., & Baumgartner, H. (2004). Multicollinearity and measurement error in structural equation models: implications for theory testing. Marketing Science, 23, 519–529.

21. Hu, L. T., & Bentler, P. M. (1998). Fit indices in covariance structure modelling: Sensitivity to underparameterized model misspecification. Psychological Methods, 3, 424–453.

22. Katz, L. F., Maliken, A. C., & Stettler, N. M. (2012). Parental meta-emotion philosophy: a review of research and theoretical framework. Child Development Perspectives, 6, 417–422.

23. Klahr, A. M., & Burt, S. A. (2014). Elucidating the etiology of individual differences in parenting: A meta-analysis of behavioral genetic research. Psychological Bulletin, 140, 544–586.

24. Kline, R. (2005). Principles and practice of structural equation modelling (methodology in the social sciences) (2^nd^ Ed.), The Guilford Press: New York.

25. Landry, S. H., Smith, K. E., Swank, P. R., & Guttentag, C. (2008). A responsive parenting intervention: the optimal timing across early childhood for impacting maternal behaviors and child outcomes. Developmental psychology, 44, 13–35.

26. Loeber, R., Green, S. M., Lahey, B. B., & Stouthamer-Loeber, M. (1991). Differences and similarities between children, mothers, and teachers as informants on disruptive child behavior. Journal of Abnormal Child Psychology, 19, 75–95.

27. Lugo-Gil, J., & Tamis-LeMonda, C. S. (2008). Family resources and parenting quality: links to children’s cognitive development across the first 3 years. Child Development, 79, 1065–1085.

28. Maccoby, E. E., Martin, J. A. Socialization in the context of the family: parent-child interaction. In: Mussen P, Hetherington E, editors. Handbook of child psychology: Vol. IV. Socialization, personality, and social development. 4th. New York: Wiley; 1983. pp. 1–101.

29. Marryat, L., Thompson, L., & Wilson, P. (2017). No evidence of whole population mental health impact of the Triple P parenting programme: findings from a routine dataset. BMC Pediatrics, 17, 40. doi:10.1186/s12887-017-0800-5.

30. Muthén, L. K., & Muthén, B. O. (2015). Mplus User’s Guide. Seventh Edition. Los Angeles, CA: Muthén & Muthén.

31. Olds, D. L., Hill, P. L., O’Brien, R., Racine, D., & Moritz, P. (2003). Taking preventive intervention to scale: the nurse-family partnership. Cognitive and Behavioral Practice, 10, 278–290.

32. Olson, S. L., Bates, J. E., Sandy, J. M., & Lanthier, R. (2000). Early developmental precursors of externalizing behavior in middle childhood and adolescence. Journal of Abnormal child Psychology, 28, 119–133.

33. Patton, G. C., Olsson, C., Bond, L., Toumbourou, J. W., Carlin, J. B., Hemphill, S. A., & Catalano, R. F. (2008). Predicting female depression across puberty: a two-nation longitudinal study. Journal of the American Academy of Child & Adolescent Psychiatry, 47, 1424–1432.

34. Podsakoff, P. M., MacKenzie, S. B., Lee, J. Y., & Podsakoff, N. P. (2003). Common method biases in behavioral research: A critical review of the literature and recommended remedies. Journal of Applied Psychology, 88, 879–903.

35. Pomerantz, A. E., & Thompson, R. A. (2008). Parents’ role in chi8ldren’s personality development: the psychological resource principle. In John O. P., & Robins, R. W., & Pervin, L. A. (Eds.), Handbook of personality: theory and research (3^rd^ ed., pp. 351–374). New York: Guilford Press.

36. Rajyaguru, P., Moran, P., Cordero, M., & Pearson, R. (2018). Disciplinary parenting practice and child mental health: evidence from the UK Millennium Cohort Study. Journal of the Americal Academy of Child & Adolescent Psychiatry, 55, S200. Doi.org/10.1016/j.jaac.2018.06.033.

37. Rudolph, K. D., Hammen, C., Burge, D., Lindberg, N., Herzberg, D., & Daley, S. E. (2000). Toward an interpersonal life-stress model of depression: The developmental context of stress generation. Development and Psychopathology, 12, 215–234.

38. Sallis, H., Smith, G. D., & Munafo, M. R. (2018). Genetics of biologically based psychological differences. Philosophical Transactions of the Royal Society, 373. doi: 10.1098/rstb.2017.0162.

39. Sessa, F. M., Avenevoli, S., Steinberg, L., & Morris, A. S. (2001). Correspondence among informants on parenting: Preschool children, mothers, and observers. Journal of Family Psychology, 15, 53.

40. Thapar, A., & McGuffin, P. (1998). Validity of the shortened Mood and Feelings Questionnaire in a community sample of children and adolescents: a preliminary research note. Psychiatry Research, 81, 259–268.

41. Yang, J., Lee, S. H., Goddard, M. E., & Visscher, P. M. (2011). GCTA: a tool for genome-wide complex trait analysis. American Journal of Human Genetics, 88, 76–82.

