## Supplemental Results and Tables for "Specific domains of early parenting, their heritability and differential association with adolescent behavioural and emotional disorders and academic achievement"

**Supplementary Material**

**Methods S1**

**Measures**

**Development of Parenting Factors**

*Process of item selection*

Potential items were extracted from questionnaires administered from pregnancy to age 5 years, including self-reports of parenting behaviour, attitudes and knowledge. All items were first categorised into dimensions according to existing theoretical parenting taxonomies (Maccoby & Martin, 1983). Items categorised as parental enjoyment, conflictual relationships, and stimulation and teaching were extracted and checked by MB & RP, resulting in approximately 50-100 items per dimension. These items were entered into separate single-factor Confirmatory Factor Analysis (CFA) models per each dimension. Three groups of items did not contribute relevant variance to early parenting dimensions: (1) items regarding expectations during pregnancy (which is different construct to actual parenting once the baby is born); (2) items relating to bedtime/ feeding and crying (as these are specific contexts and so may change general parenting); and 3) items collected from questionnaires for ages over 3 years to capture a period of time most mothers in the UK spend with their children who tend to start nursery school at age 3 years. Thus, we focused on ages 0-3 years only (13 different questionnaires) and excluded items related to expectations during pregnancy and bedtime routine.

**Analysis**

*Estimating heritability of each of the 3 parenting factors and genetic correlation between parenting factors*

Variations in different components of parenting will be driven by both genetic and environmental factors (Bornstein, 2016). Existing research has mostly relied on twin studies to estimated heritability (h^2^) of parenting components (Klahr & Burt, 2013). Heritability (h^2^) is defined as the proportion of variation in a phenotype that can be attributed to genetic differences for the particular context and timepoint, which included twin, family and adoption studies, in addition to more recent methods that use data captured by genome-wide arrays. Estimates calculated according to these family-based models can be thought of as ‘true’ h^2^. These estimates will incorporate h^2^ due to variants across the entire genome, including rare variants and those not captured by single nucleotide polymorphisms (SNPs) included on genotyping platforms (Yang, Lee, Goddard, & Visscher, 2011). However, twin models, by design, use closely related individuals. These individuals are thus assumed to share a great deal of their environment, however, if the equal environments assumption is violated this can impact on h^2^ estimates. For example, more similar treatment of identical (MZ) twins could lead to false inflation of estimates (Sallis, Davey Smith, & Munafo, 2017). Here we describe different components of parenting experiences and estimate SNP based h^2^ based on maternal molecular genetic data. This has not been applied to the heritability of parenting before.

#### First, estimates of SNP-based heritability (h^2^_SNP_) for each parenting factor were calculated using the restricted maximum likelihood (REML) method implemented within the genome-wide complex trait analysis (GCTA) software (Yang et al., 2011). Second, we used a bivariate REML approach to estimate the genetic correlation between each of the parenting factors with each other to investigate shared genetic architecture across each of the parenting factors. Any overlap here could be due to pleiotropy (genetic effects on multiple traits), shared biological mechanisms between domains, or a causal relation from one domain to another.

*Latent factor model*

Once items that were both theoretically sound and showed standardised loadings >0.15 on the relevant dimension were established, we developed a combined model using Confirmatory Factor Analyses (CFA) with a robust weighted least square (WLSMV) estimator in Mplus, which provides the best option to model categorical data (Brown, 2012). We then loaded items to the hypothesised dimension and compared modification indices to highlight items which may be captured better by a different factor. Similar items that were collected at different time points were set to correlate with each other in the model to account for shared variance related to time and the repeated nature of questions. The RMSEA (0.024, 95% CI 0.024 to 0.025) and the CFI (0.92) indicated that the measurement model fit the data well, supporting the adequacy of the model for tests of structural paths. The flow chart of items included into the CFA is presented in Figure S1. Derived factors, items and factor loadings are presented in Table S1. There were relatively high correlations between parenting factors (Figure 1).

**Results S1**

*Associations between parenting factor scores and confounding variables*

Overall, there were some differences in parenting according to child gender. Mothers reported less parental enjoyment and stimulation, and more conflictual relationships when parenting boys. Mothers with higher educational attainment (non-compulsory secondary level (up to age 18 years) and university level education) reported more enjoyment and less conflictual relationships and stimulation and teaching activities with their child than mothers with lower educational attainment (compulsory secondary level (up to age 16 years), minimal education or none). Younger mothers (≤20 years old) reported more enjoyment, stimulation and teaching activities, and conflictual relationships with their child than older mothers (≥20 years old). Mothers who experienced depression during pregnancy reported more conflictual relationships, less parental enjoyment and stimulation activities with the child than mothers with no depression.

*Associations between parenting factors and offspring behavioural disorders, depressive symptoms and academic achievement at 16 years*

We found evidence for an interaction between maternal conflictual relationships and enjoyment with offspring behavioural disorders at age 16 years (interaction term β=0.113, p<0.001). Figure S2 represents percentage of offspring with conduct disorder (CD) or oppositional defiant disorder (ODD) diagnosis according to patterns of parental conflict and enjoyment. In our sample 6.2% of the offspring who experienced early parent-child conflict were diagnosed with ODD and CD, which is almost double the population prevalence of 3.4% (Egger & Angold, 2006). However, if conflict occurred in the context of high parental enjoyment, only 3.5% of offspring were diagnosed with ODD and CD.

Characteristics of the sample by the completeness of the data (Table S3) suggested that mothers in the study sample with complete parenting data were more likely to be older, have one child, and to report higher educational attainment, lack of financial problems, and less likelihood of antenatal depression when compared to the original Avon Longitudinal Study of Parents and Children (ALSPAC) cohort. Given these differences, we repeated our analyses using complete case sample on all variables (n=2,694) with comparable results. Specifically, there was evidence for an association between conflictual relationships and offspring behavioural disorders at age 16 years in the fully adjusted model (β=0.269, p<0.001). There was also evidence for an association between stimulating and teaching activities and grades in English language (β=0.081, p=0.003).

**Figure S1.** Flow chart depicting items included into the final Confirmatory Factor Analysis (CFA) model

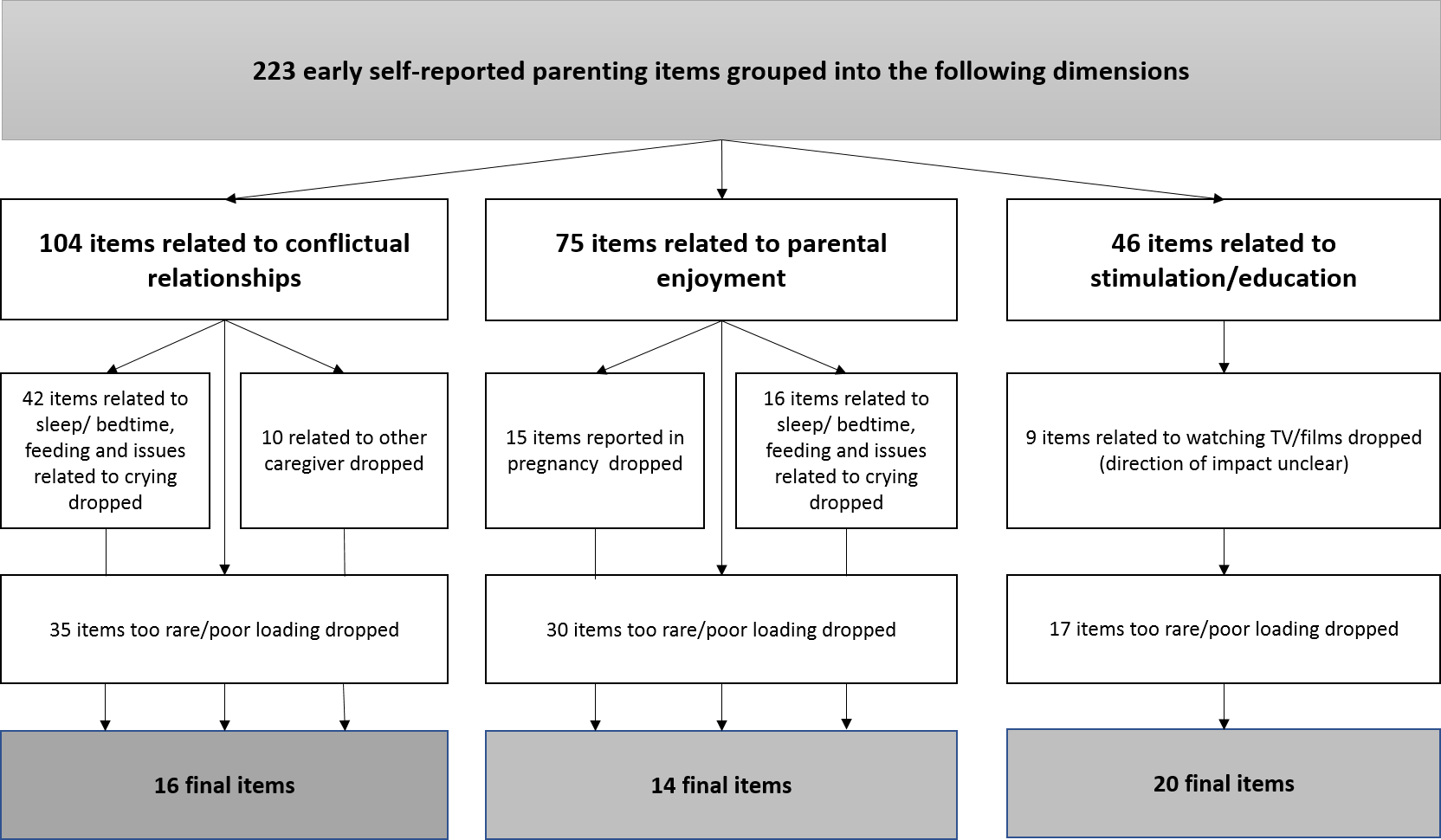

**Figure S2.** Percentage of offspring diagnosed with behavioural disorder (CD or ODD) according to patterns of maternal conflict and enjoyment.

|  | % with behaviour disorder (CD or ODD) |
| --- | --- |
| High conflict-High enjoy | 3.5 |
| High Conflict | 5.9 |
| High conflict-Low enjoy | 6.65 |
| Low enjoy | 5.4 |
| Low conflict Low enjoy | 2.5 |

*Note:* High or low levels refer to mothers being in the top or bottom tertile of the relevant parenting factor score.

**Table** **S1.** Derived factors, items, standardised factor loadings and fit indices for the measurement model.

| Age/Factor | Items | Factor loading | S.E. |
| --- | --- | --- | --- |
| **Conflictual relationship** |  |  |  |
| 1 year 6 months | Do you ever have a battle of wills with your toddler? | 0.343 | 0.013 |
| 1 year 6 months | When she has temper tantrums how often do you smack? | 0.266 | 0.014 |
| 1 year 6 months | When she has temper tantrums how often do you shout? | 0.337 | 0.014 |
| 2 years 6 months | Do you ever have a battle of wills with your child? | 0.431 | 0.013 |
| 2 years 6 months | Who most often wins? | 0.149 | 0.015 |
| 3 years 6 months | Do you feel that your child's crying is a problem? | 0.224 | 0.014 |
| 3 years 6 months | Do you ever have a battle of wills with your child? | 0.453 | 0.014 |
| 3 years 6 months | Who most often wins? | 0.148 | 0.016 |
| 3 years 6 months | When he has temper tantrums how often do you leave your child? | 0.261 | 0.016 |
| 3 years 6 months | When he has temper tantrums how often do you ignore your child? | 0.291 | 0.015 |
| 3 years 6 months | When he has temper tantrums how often do you smack your child? | 0.335 | 0.015 |
| 3 years 6 months | When he has temper tantrums how often do you shout at your child? | 0.450 | 0.014 |
| 3 years 6 months | When he has temper tantrums how often do you tell your child off? | 0.363 | 0.015 |
| 3 years 6 months | When he has temper tantrums how often do you bribe your child? | 0.451 | 0.014 |
| 3 years 11 months | This child gets on my nerves | 0.447 | 0.013 |
| 3 years 11 months | Mother feels very unattached to child | 0.296 | 0.014 |
| **Parental enjoyment** |  |  |  |
| 4 weeks | Often mothers are surprised how long it takes to love their babies. How long has it taken you? | 0.272 | 0.011 |
| 8 months | I really enjoy my baby | 0.587 | 0.009 |
| 8 months | It is a great pleasure to watch my baby develop | 0.391 | 0.011 |
| 8 months | Having a baby has made me feel more fulfilled | 0.522 | 0.010 |
| 8 months | Babies are fun | 0.625 | 0.008 |
| 1 year 9 months | Toddlers are fun | 0.620 | 0.009 |
| 1 year 9 months | I really love my toddler | 0.366 | 0.011 |
| 1 year 9 months | It is a great pleasure to watch my child grow | 0.448 | 0.010 |
| 1 year 9 months | My child gives me great joy | 0.558 | 0.009 |
| 2 year 9 months | I really enjoy this child | 0.588 | 0.009 |
| 2 year 9 months | It is a great pleasure to watch my child develop | 0.451 | 0.011 |
| 2 year 9 months | Having this child has made me feel more fulfilled | 0.570 | 0.010 |
| 2 year 9 months | Children are fun | 0.664 | 0.008 |
| 3 years 6 months | How often do you cuddle your son/daughter? | 0.048 | 0.012 |
| **Stimulation and teaching** |  |  |  |
| 6 months | Do you try to teach your child? | 0.461 | 0.013 |
| 6 months | Do you talk to your baby while you work (e.g., while you do housework)? | 0.412 | 0.013 |
| 6 months | How often do you do these activities with your baby (e.g., play with toys)? | 0.319 | 0.014 |
| 6 months | How often do you do these activities with your baby (e.g., show her pictures in books)? | 0.508 | 0.012 |
| 6 months | How often do you do these activities with your baby (e.g., physical play such as clapping, rolling over)? | 0.341 | 0.013 |
| 6 months | How often do you play with your baby? | 0.323 | 0.013 |
| 6 months | About how often do you take her to places of entertainment? | 0.170 | 0.012 |
| 1 year 6 months | Which things do you try to do with her (e.g., parts of the body)? | 0.198 | 0.013 |
| 1 year 6 months | Which things do you try to do with her (e.g., colours)? | 0.403 | 0.012 |
| 1 year 6 months | Which things do you try to do with her (e.g., alphabet) | 0.334 | 0.012 |
| 1 year 6 months | Which things do you try to do with her (e.g., numbers)? | 0.377 | 0.012 |
| 1 year 6 months | Which things do you try to do with her (e.g., nursery rhymes)? | 0.193 | 0.013 |
| 1 year 6 months | Which things do you try to do with her (e.g., songs)? | 0.189 | 0.013 |
| 1 year 6 months | Which things do you try to do with her (e.g., shapes and sizes)? | 0.352 | 0.012 |
| 1 year 6 months | Which things do you try to do with her (e.g., politeness)? | 0.138 | 0.013 |
| 2 years | When you are at home with your child, how often do you do the following: go out to a park or playground with him? | 0.266 | 0.012 |
| 2 years 6 months | About how often do you take him to park or playground? | 0.239 | 0.013 |
| 2 years 6 months | Do you try to teach your child? | 0.466 | 0.011 |
| 3 years 6 months | About how often is he taken to local shops? | 0.084 | 0.013 |
| 3 years 6 months | About how often is he taken to park or playground? | 0.207 | 0.013 |
| **Model Fit Indices** | | | |
| Free parameters | | 315 | |
| Root Mean Square Error of Approximation (RMSEA) | | 0.024 (95%CI 0.024 to 0.025) | |
| Comparative Fit Index (CFI) | | 0.918 | |
| Tucker-Lewis Index (TLI) | | 0.902 | |

**Table S2.** Associations between parenting factor scores (enjoyment, conflictual relationships, stimulation/teaching) and child and parental confounders

| Parenting factors | Child gender | | Maternal educational attainment | | Maternal age | | Maternal depression | |
| --- | --- | --- | --- | --- | --- | --- | --- | --- |
|  | Male  (n=5,725) | Female  (n=5,388) | Degree/  A Level  (n=6,131) | O Level/  None  (n=3,994) | ≤20 years  (n=777) | ≥20 years  (n=10,336) | No  (n=7,727) | Yes  (n=2,085) |
| Enjoyment,  *Mean (SD)* | 0.007  (0.268) | -0.007  (0.260) | -0.008  (0.264) | 0.019  (0.271) | -0.019  (0.248) | 0.001  (0.266) | -0.017  (0.252) | 0.075  (0.307) |
| ANOVA t-test, p-value | 2.975, 0.003 | | -4.873, <0.001 | | 19.91, <0.001 | | -14.06, <0.001 | |
| Conflictual relationships,  *Mean (SD)* | 0.008  (0.261) | -0.009  (0.261) | -0.005  (0.264) | 0.016  (0.259) | 0.015  (0.265) | -0.001  (0.261) | -0.018  (0.253) | 0.080  (0.283) |
| ANOVA t-test, p-value | 3.501, <0.001 | | -3.866, <0.001 | | 1.657, 0.098 | | -15.444, <0.001 | |
| Stimulation/Teaching,  *Mean (SD)* | 0.004 (0.161) | -0.005 (0.156) | 0.006  (0.159) | -0.015  (0.156) | -0.018  (0.149) | 0.001  (0.159) | -0.004  (0.157) | 0.017  (0.165) |
| ANOVA t-test, p-value | 3.339, 0.001 | | 6.731, <0.001 | | -3.269, 0.001 | | -9.13, <0.001 | |

*Note:* p-values based on ANOVA t-test for differences in means for continuous factor scores. Enjoyment: higher scores represent lower parental enjoyment; Conflictual relationships: higher scores represent higher levels of conflict; Stimulation/Teaching: higher scores represent lower levels of stimulation.

**Table S3.** Distribution of key sociodemographic characteristics in the original Avon Longitudinal Study of Parents and Children (ALSPAC) cohort and the study sample with complete parenting data

| Sociodemographic characteristics | Core ALSPAC sample | Study sample with complete parenting data |
| --- | --- | --- |
|  | *n (%)* | *n (%)* |
| *Maternal educational attainment* | | |
| Degree/A Level | 84 (24.1%) | 4,328 (38.1%) |
| O Level/None | 265 (75.9%) | 7,043 (61.9%) |
| Chi^2^, p-value | 28.25, <0.001 | |
| *Maternal age, mean (SD)* | 26.6 (5.6) | 27.4 (4.9) |
| β, p-value | 0.82, <0.001 | |
| *Maternal antenatal depression* | | |
| No | 647 (79.8%) | 9,826 (86.6%) |
| Yes | 164 (20.2%) | 1,526 (13.4%) |
| Chi^2^, p-value | 29.08, <0.001 | |
| Parity | | |
| 1 child | 675 (75.3%) | 9,787 (80.0%) |
| >1 child | 221 (24.7%) | 2,443 (20.0%) |
| Financial problems |  |  |
| No | 658 (72.0%) | 9,782 (78.6%) |
| Yes | 255 (28.0%) | 2,666 (21.4%) |

*Note:* p-values based on Chi^2^ test of the association between completeness of the sample and categorical variables, and linear regression β coefficients for differences in means for continuous variables; sample sizes vary due to the differences in data availability on sociodemographic characteristics

**Table S4.** Standardised path coefficients of the associations between parenting factors and offspring behavioural disorders, depressive symptoms and academic achievement at 16 years (n=12,358)

| Offspring outcomes | Model estimates | | | |
| --- | --- | --- | --- | --- |
|  | β | SE | p-value | 95% CI |
| *Behavioural disorders (DAWBA)* | | | | |
| Parental enjoyment | 0.093 | 0.062 | 0.132 | -0.029 to 0.215 |
| Conflictual relationships | 0.227 | 0.084 | 0.007 | 0.062 to 0.391 |
| Stimulation/teaching | -0.042 | 0.051 | 0.416 | -0.142 to 0.058 |
| *Depressed mood (SMFQ)* | | | | |
| Parental enjoyment | 0.015 | 0.032 | 0.651 | -0.048 to 0.078 |
| Conflictual relationships | -0.063 | 0.044 | 0.151 | -0.149 to 0.023 |
| Stimulation/teaching | 0.013 | 0.026 | 0.624 | -0.038 to 0.064 |
| *Academic achievement (GCSE in English)* | | | | |
| Parental enjoyment | 0.015 | 0.032 | 0.651 | -0.048 to 0.078 |
| Conflictual relationships | -0.063 | 0.044 | 0.151 | -0.149 to 0.023 |
| Stimulation/teaching | 0.013 | 0.026 | 0.624 | -0.038 to 0.064 |

*Note:* analyses adjusted for parental (maternal age, educational attainment, depression) and child (gender and early behavioural problems) confounders.

**References**

1. Maccoby, E. E., Martin, J. A. Socialization in the context of the family: Parent-child interaction. In: Mussen P, Hetherington E, editors. Handbook of child psychology: Vol. IV. Socialization, personality, and social development. 4th. New York: Wiley; 1983. pp. 1–101.

2. Bornstein, M. H. Determinants of parenting. (2016). In D. Cicchetti (Ed.), Developmental Psychopathology: Risk, Resilience, and Intervention (3rd ed., Vol. 4, pp. 180-270). Hoboken, NJ: Wiley.

3. Brown, T. A., & Moore, M. T. (2012). Confirmatory factor analysis. *Handbook of structural equation modeling*, 361-379.
4. Egger, H. L., & Angold, A. (2006). Common emotional and behavioral disorders in preschool children: presentation, nosology, and epidemiology. *Journal of Child Psychology and Psychiatry*, *47*, 313-337.

5. Klahr, A. M., & Burt, S. A. (2014). Elucidating the etiology of individual differences in parenting: A meta-analysis of behavioral genetic research. *Psychological Bulletin*, *140*, 544-586.

6. Sallis, H., Smith, G. D., & Munafo, M. R. (2018). Genetics of biologically based psychological differences. *Philosophical Transactions of the Royal Society, 373*. doi: 10.1098/rstb.2017.0162.

7. Yang, J., Lee, S. H., Goddard, M. E., & Visscher, P. M. (2011). GCTA: a tool for genome-wide complex trait analysis. *American Journal of Human Genetics, 88*, 76-82.
